# Protein interactions between the oral bacteria *Fusobacterium nucleatum* and *Porphyromonas gingivalis* in biofilm and planktonic culture

**DOI:** 10.1101/2020.11.23.393850

**Authors:** Marwan Mansoor Ali Mohammed, Veronika Kuchařová Pettersen, Audun H. Nerland, Harald G. Wiker, Vidar Bakken

## Abstract

The opportunistic pathogens *Fusobacterium nucleatum* and *Porphyromonas gingivalis* are Gram-negative bacteria associated with oral biofilm and periodontal disease. Although liquid cultures are often the preferred cultivation method in microbiology, bacterial cells in biofilm adopt a profoundly different phenotype reflecting the close cell-to-cell contact compare to their planktonic counterparts. To investigate *F. nucleatum* and *P. gingivalis* interactions relevant in biofilm formation, we applied liquid chromatography-tandem mass spectrometry to determine the expressed proteome of *F. nucleatum* and *P. gingivalis* cells that were grown either as biofilm or in planktonic culture, and individually or together.

The proteomic analyses detected 1,322 *F. nucleatum* and 966 *P. gingivalis* proteins. We statistically compared the proteins label-free quantitative (LFQ) intensities between biofilm and planktonic culture and identified significant changes (p-value ≤0.05) in 0,4% *F. nucleatum* proteins, 7% *P. gingivalis* proteins, and more than 14% of all proteins in the dual-species model. For both species, proteins involved in vitamin B2 (riboflavin) metabolic process had significantly increased levels in the biofilm condition. In both mono- and dual-species biofilm models, *P. gingivalis* increased the production of proteins functional in translation, oxidation-reduction, and amino acid metabolism, when compared to planktonic cultures. However, when we compared LFQ intensities between mono- and dual-species models, over 90% of the significantly changed *P. gingivalis* proteins had their levels reduced in biofilm and planktonic settings of the dual-species model.

Our findings suggest that the two bacteria interact with each other at the protein level and indicate that *P. gingivalis* reduces the production of multiple proteins because of more favourable growth conditions provided by *F. nucleatum* presence. The results highlight the complex interactions of bacteria contributing to oral biofilm, which need to be considered in the design of future prevention strategies.

## Introduction

*Fusobacterium nucleatum* and *Porphyromonas gingivalis* are important colonizers of the subgingival biofilms (1). Both bacteria play a role in the pathogenesis of periodontal diseases, a group of inflammatory diseases of the teeth supporting tissues (2). A mild form called gingivitis is highly prevalent and can affect up to 90% of the worldwide population. However, gingivitis does not affect the underlying supporting structures of the teeth and is reversible. A more severe form of the disease, periodontitis, results in loss of connective tissue and bone support and is the main cause of tooth loss in adults (3).

*F. nucleatum* is commonly cultivated from the subgingival biofilm and tends to aggregate with other oral bacteria, working as a bridge between early and late colonizers in the development of the dental biofilm (4). *P. gingivalis* is a member of the Socransky’s red complex, a group of bacteria strongly associated with periodontal disease (5). It is often found in the deep periodontal pockets, and it produces a broad array of potential virulence factors involved in tissue colonization and destruction as well as in perturbations of the host defence (reviewed in (6–8)).

*P. gingivalis* and *F. nucleatum* are considered strict anaerobes, and both species display a synergistic enhancement in biofilm formation and pathogenicity (9, 10). Several studies showed that the bacteria could grow in a partially oxygenated condition when grown together and suggested that enhanced production of oxidoreductive enzymes by *F. nucleatum* is protecting *P. gingivalis* from oxidative stress (11, 12). Similarly, *in vitro* and *in vivo* models showed a nearby association between the bacteria, indicating that they co-aggregate and potentially support each other (10, 13, 14). However, how the bacteria interact at the protein level remains poorly understood.

Although most microbiology research has been focused on free-floating bacteria in suspension (planktonic cells), mounting evidence indicates that cells growing in biofilms are in a very different physiological state (15–17). For example, the envelope fraction of *Pseudomonas aeruginosa* cells grown as a biofilm showed a 30–40% difference in the detected proteins compared with the same fraction of *P. aeruginosa* planktonic cells (15). Recent investigations indicate that biofilm is the preferred form of life for most microbes, particularly those of pathogenic nature (18).

In the current study, we explored the differences between cells grown in a culture or biofilm at the proteome level. To address the synergistic relationship between *F. nucleatum* and *P. gingivalis*, we grew the bacteria both individually and in a dual-species model. The results suggest that the two bacteria actively interact with each, and *P. gingivalis* reduces the production of certain proteins in the biofilm, possibly because of a favourable presence of *F. nucleatum* proteins that support the necessary biochemical adaptations.

## MATERIALS AND METHODS

### Bacterial strains and growth conditions

*Fusobacterium nucleatum* subsp. *nucleatum* type strain ATCC 25586 and *Porphyromonas gingivalis* type strain ATCC 33277 were used in the current study. Anaerobic conditions (5% CO_2_, 10% H_2_, and 85% N_2_) (Anoxomat System, MART Microbiology, Lichtenvoorde, The Netherlands) were used in all steps of the experiment. The bacterial strains were grown on fastidious anaerobic agar (FAA) plates at 37°C for 48 hours. Few colonies were then inoculated in Brucella broth (Becton Dickinson, Maryland, USA) supplemented with five μg/ml hemin and 0.25 μg/ml Vitamin K. The bacteria were grown overnight in the liquid medium at 37°C. The overnight cultures were adjusted to an absorbance of 0.15 at 600 nm (A_600_), of which 10 ml (5 ml from each species in dual-species biofilm) was transferred to a separate 25 cm^2^ (area) polystyrene cell culture flasks (TPP Techno Plastic Products, Trasadingen, Switzerland). The flasks were incubated at 37°C for four days. The medium was then removed and the biofilm samples were washed once with phosphate buffered saline (PBS), before the biofilms were harvested with cell scraper (Nunc, Rochester, NY, USA). The biofilm samples were resuspended in 500 μl PBS and stored at −20°C until further processing.

The planktonic cultures were anaerobically grown in the same liquid medium as described above in 10 ml glass round bottom test tubes with screw caps at 37°C, without shaking. After four days, the bacteria were collected by centrifugation at 3000 x *g* for 3 min in room temperature. The pelleted cells were resuspended in 500 μl PBS and stored at −20°C until further processing.

### Protein extraction from the biofilm and planktonic samples

The bacterial cells were washed 3 times by resuspension in 1 ml PBS and centrifugation for 10 min each time at 6000 x *g* at +4°C. In a final step, the cells were resuspended in 1 ml of extraction buffer (10 mM Tris-HCl, 2.5 % SDS, pH 8.0). The cell suspensions were transferred to FastPrep® Lysing Matrix A, 2 mL Tube (MP Biomedicals, California, USA) and then bead-beated in FastPrep® FP120 Cell Homogenizer (Thermo, California, USA) for 45 sec at 6.5 m/s. The cell extracts were cooled on ice for 5 min, then the cell debris was removed by 30 min centrifugation at 10,000 x *g*, +4°C. The collected supernatant was kept on ice until measurements of protein concentrations using Direct Detect® Spectrometer (Merck Millipore, Darmstadt, Germany).

### Sample preparation for the proteomic analysis

Protein extracts from the biofilm and planktonic cells prepared from three independent biological replicates were subjected to the Filter Aided Sample Preparation method (19). The protein samples were mixed with a solution of 10 mM dithiothreitol (DTT) in 100 mM ammonium bicarbonate (NH_4_HCO_3_) [solution to total protein ratio (v/w) 1:10] and incubated for 45 min at 56°C. The Microcon device YM-10 filters (Merck Millipore, Darmstadt, Germany) were first conditioned by adding 100 μl of urea buffer (8 M urea, 10 mM HEPES, pH 8.0) and centrifuged at 14,000 x *g* for 5 minutes. Aliquots of the samples containing 50 μg of protein were mixed with 200 μl urea buffer in the filter unit and centrifuged at 14,000 x *g* for 15 min, and this step was repeated. The filtrate was discarded, and 100 μl of 0.05 M iodoacetamide was added to each sample. The samples were mixed at 600 rpm for 1 min and incubated without mixing in the dark for 20 min, followed by centrifugation at 14,000 x *g* for 10 min, three washes with 100 μl urea buffer, and three washes with 100 μl 40 mM NH_4_HCO_3_. Proteins retained on the filter were digested with trypsin (Thermo Fisher Scientific, IL, USA) in 40 mM NH_4_HCO_3_ buffer (enzyme to protein ratio 1:50) at 37°C for 16 h. The released peptides were collected by adding 50 μl of MS grade water followed by centrifugation at 14,000 x *g* for 15 min. This step was repeated twice. The samples were concentrated in a vacuum concentrator (Eppendorf, Hamburg, Germany).

### Filtration and desalting

StageTips to be used for filtration and peptide samples desalting were prepared in-house according to the protocol developed by Rappsilber and colleagues (20). Shortly, 3M Empore C18 extraction disks (3M, Minnesota, USA) were packed in 200 μl pipet tips by a blunt-ended needle and a plunger or metal rod that helped fit the extracted disks in the pipet tips. The disks were then wetted by passing 20 μl of methanol, followed by 20 μl of elution buffer [80% acetonitrile (ACN), 0.1% formic acid (FA)]. The disks were conditioned and equilibrated with 20 μl of 0.1% FA just before the last residue of the previous buffer left the tip to avoid drying of the disks. The prepared peptide mixtures (volumes 20-40 μl) were loaded on top of the Stage Tip. The peptide samples were first desalted by washing with 20 μl of 0.1% FA and then eluted by adding 20 μl of the elution buffer two times. The collected samples were dried in the vacuum concentrator and stored at −80°C until further analyses. Peptide samples were resuspended by adding 1 μl of 100% FA and 19 μl of 2% ACN prior to liquid chromatography-tandem mass spectrometry (LC-MS/MS) analysis.

### LC-MS/MS

The MS/MS analysis was carried out at the Proteomics Unit, University of Bergen (PROBE), on an Ultimate 3000 RSLC system (Thermo Scientific) connected to a linear quadrupole ion trap-Orbitrap (LTQ-Orbitrap) MS (Thermo Scientific) equipped with a nanoelectrospray ion source. Briefly, 1 μg protein was loaded onto a pre-concentration column (Acclaim PepMap 100, 2 cm ×75 μm i.d. nanoViper column, packed with 3 μm C18 beads) at a flow rate of 5 μl/min for 5 min using an isocratic flow of 0.1% trifluoroacetic acid, vol/vol (TFA). Peptides were separated during a biphasic ACN gradient from two nanoflow ultra-performance liquid chromatography (UPLC) pumps (flow rate of 270 nl/min) on the analytical column (Acclaim PepMap 100, 50cm x 75μm i.d. nanoViper column, packed with 3μm C18 beads). Solvent A and B was 0.1% FA (vol/vol) in water or ACN (vol/vol), respectively. Separated peptides were sprayed directly into the MS instrument during a 195 min LC run with the following gradient composition: 0-5 min 5% B, 5-6 min 8% B, 6-135 min 7–32% B, 135-145 min 33– 40% B, and 145-150 min 40-90% B. Elution of very hydrophobic peptides and conditioning of the column was performed by isocratic elution with 90% B (150-170 min) and 5% B (175-195 min), respectively. Desolvation and charge production were accomplished by a nanospray Flex ion source.

The mass spectrometer was operated in the data-dependent-acquisition mode to automatically switch between Orbitrap-MS and LTQ-MS/MS acquisition. Survey of full-scan MS spectra (from m/z 300 to 2,000) were acquired in the Orbitrap with resolution of R = 240,000 at m/z 400 (after accumulation to a target of 1,000,000 charges in the LTQ). The method used allowed sequential isolation of the most intense ions (up to 10, depending on signal intensity) for fragmentation on the linear ion trap using collision-induced dissociation at a target value of 10,000 charges. Target ions already selected for MS/MS were dynamically excluded for 18s. General mass spectrometry conditions were as follows: electrospray voltage, 1.8 kV; no sheath; and auxiliary gas flow. Ion selection threshold was 1,000 counts for MS/MS, and an activation Q-value of 0.25 and activation time of 10 ms was also applied for MS/MS.

### Data analysis

The acquired MS raw data were processed by using the MaxQuant software (21), version 1.5.2.8, with default settings and the following additional options: Label-Free Quantification (LFQ) (22), match between runs, and 0.01 false discovery rate (FDR) at both peptide and protein level. The MS spectra were searched against protein databases of *F. nucleatum* type strain ATCC 25586 and *P. gingivalis* type strain ATCC 33277, downloaded from the Universal Protein Knowledgebase on the 4^th^ of February 2015. Normalized spectral proteins intensities (LFQ intensity), which are proportional to the quantity of a given protein in a sample, were derived by the MaxLFQ algorithms (22). The normalization ensures that one can compare LFQ scores between different samples.

MaxQuant output data were analysed with the Perseus module (23). The post MaxQuant analysis included filtering of the generated ‘proteingroups.txt’ table for contaminants, only identified by site and reverse hits. Each protein identified in at least two of the three replicates was considered valid. Proteins with significant differential levels were identified by statistical analysis based on two-sided t-test, which was performed on proteins log_2_ transformed LFQ values. A protein was considered significantly changed if it was marked as significant in the t-test and showed more than 2 log_2_ difference form the the mean LFQ intensity.

Functional protein classification was performed by using The Database for Annotation, Visualization and Integrated Discovery (DAVID) (24) and QuickGO annotation database (25). Potentially interesting clusters identified by DAVID were studied individually. The web-based application SOSUI-GramN (26) was used to predict the subcellular localization of the identified proteins. The mass spectrometry proteomics data have been deposited to the ProteomeXchange Consortium via the PRIDE partner repository with the dataset identifier PXD00828 (Username, Password: R3hEWdEI) (27, 28).

## Results and Discussion

This study’s objective was to investigate *F. nucleatum* and *P. gingivalis* proteins relevant in biofilm formation. We grew the bacteria in mono- or dual-species model either as biofilms or planktonic cells. Growing the bacteria both individually and together allowed us to investigate interactions between the two species at the protein level. Altogether, six different growth conditions (biofilm and planktonic culture of *F. nucleatum*, *P. gingivalis,* and the dual-species model) were analysed by LC-MS/MS using three biological replicates for each condition.

The analysis yielded approximately a million MS/MS spectra, which we searched against protein databases of either *F. nucleatum* or *P. gingivalis*. The data search matched the spectra to 23,423 distinct peptide sequences (**Table S1**, Supplementary file 1), which were assigned to 2,288 different proteins (**Table S2**, Supplementary file 1). The number of identified proteins in each growth condition and their predicted subcellular localization are shown in **Table 1**.

**Table 1.** The number of identified proteins in each condition categorized according to predicted cellular localization.

| Subcellular localization<sup>#</sup> | Biofilm |  |  | Planktonic culture |  |  |
| --- | --- | --- | --- | --- | --- | --- |
|  | FnBio | PgBio | FnPgBio | FnPla | PgPla | FnPgPla |
| **Cytoplasmic** | 893 | 552 | 998 | 904 | 483 | 1132 |
| **Extracellular** | 45 | 27 | 56 | 44 | 28 | 62 |
| **Inner membrane** | 210 | 131 | 216 | 216 | 125 | 267 |
| **Outer membrane** | 34 | 92 | 95 | 36 | 88 | 107 |
| **Periplasm** | 21 | 73 | 69 | 20 | 72 | 72 |
| **Unknown** | 54 | 42 | 58 | 54 | 38 | 78 |
| **Total** | 1257 | 917 | 1492 | 1274 | 834 | 1718 |
| **Proteome coverage\* (%)** | 61.3 | 45.3 | 36.6 | 62.2 | 41.2 | 42.2 |

Abbreviations: Fn = *F. nucleatum*, Pg = *P. gingivalis*, Bio = Biofilm, Pla = Planktonic
<sup>#</sup> The predictions were made by using the web-based application SOSUI-GramN (26).
\* The number of detected proteins divided by the theoretical proteome (Fn 2049 proteins, Pg 2022 proteins) derived from the UniProt database.

The proteome coverage, which we defined as the number of the detected proteins divided by the theoretical proteome derived from the UniProt database, was higher for *F. nucleatum* (≈ 62%) than the one of *P.* gingivalis (≈ 43%), both in biofilm and planktonic conditions (**Table 1**). The numbers of detected proteins were similar to previous studies that reported proteome coverage between 48 and 60% for *P. gingivalis* (29, 30) and 58% for *F. nucleatum* (31).

Up to 84% of the identified proteins (1,916) were described by LFQ intensities, which indicate relative protein levels in the analysed samples (**Table S3**, Supplementary file 1). The LFQ intensities covered a dynamic range of ≈ 12 log_2_ (**Fig S1**, Supplementary file 2), and the correlations between replicates, represented as the Pearson correlation coefficient, varied between 0.79-0.98 (**Fig S2**, Supplementary file 2).

The most abundant proteins identified in the biofilm (**Table 2**) and planktonic lifestyles (**Table 3**) included oxidoreductases, acyltransferases, outer membrane proteins, and proteases, among others. We identified major virulence factors of *F. nucleatum* and *P. gingivalis* (*e.g*., major outer membrane protein FomA and Lys-gingipain) as well as cytoplasmic proteins like Acetyl-CoA acetyltransferase, Alkyl hydroperoxide reductase C22 protein, and Neutrophil-activating protein A, which were previously shown to be abundant in the biofilms extracellular polymeric matrix (32). The latter finding confirms that proteins released from dead cells are used in the biofilm matrix (33).

**Table 2.** Top ten abundant proteins identified in the mono- and dual-species biofilms of *F. nucleatum* and *P. gingivalis.*

| UniProt<br>Accession | Gene Name | Protein Description | Major Function | Log<sub>2</sub> LFQ |
| --- | --- | --- | --- | --- |
| Top 10 in *F. nucleatum* biofilm |  |  |  |  |
| Q8RG30 | *FN0488* | Glutamate dehydrogenase | oxidoreductase | 32.66 |
| Q8RES0 | *FN1024* | DNA-binding protein HU | DNA binding protein | 32.08 |
| Q8RIN6 | *FN1535* | Acyl-CoA dehydrogenase, short-chain specific | oxidoreductase | 32.02 |
| Q8R6D3 | *FN1983* | Alkyl hydroperoxide reductase C22 protein | oxidoreductase | 31.90 |
| Q8R643 | *FN1170* | Pyruvate-flavodoxin oxidoreductase | oxidoreductase | 31.79 |
| Q8REE2 | *FN1165* | D-galactose-binding protein | metal ion binding protein | 31.72 |
| Q8RHQ6 | *FN1943* | Tryptophanase | lyase | 31.57 |
| Q8RIP5 | *FN1526* | Fusobacterium outer membrane protein family | OMP/adhesin | 31.43 |
| Q8RHY1 | *FN1859* | Major outer membrane protein FomA | OMP/VF | 31.29 |
| Q8RG24 | *FN0495* | Acetyl-CoA acetyltransferase | acetyltransferase | 31.28 |
| Top 10 in *P. gingivalis* biofilm |  |  |  |  |
| B2RM93 | *rgpA* | Arginine-specific cysteine proteinase RgpA | protease/VF | 32.04 |
| B2RL46 | *rplL* | 50S ribosomal protein L7/L12 | ribosomal protein | 31.17 |
| B2RLK2 | *kgp* | Lys-gingipain | protease/VF | 31.04 |
| B2RKJ1 | *gdh* | NAD-specific glutamate dehydrogenase | oxidoreductase | 31.03 |
| B2RLK7 | *hagA* | Hemagglutinin protein HagA | agglutinate erythrocytes/ VF | 30.85 |
| B2RHG7 | *ragA* | Receptor antigen A | protein with receptor activity | 30.66 |
| B2RHG8 | *ragB* | Receptor antigen B | protein with receptor activity | 30.52 |
| B2RHA9 | *PGN_0235* | DNA-binding protein HU | DNA binding protein | 30.52 |
| B2RIQ1 | *PGN_0727* | 4-hydroxybutyryl-CoA dehydratase | oxidoreductase | 30.03 |
| B2RII3 | *PGN_0659* | 35 kDa hemin binding protein | hemin binding protein /VF | 29.44 |
| Top 10 in dual-species biofilm |  |  |  |  |
| B2RM93 | *rgpA* | Arginine-specific cysteine proteinase RgpA | protease/VF | 32.75 |
| B2RLK2 | *kgp* | Lys-gingipain | protease/VF | 32.59 |
| Q8RG24 | *FN0495* | Acetyl-CoA acetyltransferase | acetyltransferase | 32.11 |
| Q8RES5 | *FN1019* | 3-hydroxybutyryl-CoA dehydrogenase | oxidoreductase | 31.74 |
| B2RLK7 | *hagA* | Hemagglutinin protein HagA | agglutinate erythrocytes/ VF | 31.47 |
| B2RHG7 | *ragA* | Receptor antigen A | protein with receptor activity | 31.30 |
| Q8RFC2 | *hutH1* | Histidine ammonia-lyase 1 | lyase | 31.12 |
| Q8RHY1 | *FN1859* | Major outer membrane protein FomA | OMP/ VF | 31.02 |
| Q8R6D3 | *FN1983* | Alkyl hydroperoxide reductase C22 protein | oxidoreductase | 30.99 |
| Q8RGB0 | *FN0396* | Dipeptide-binding protein | peptide transporter | 30.88 |

Abbreviations: OMP = outer membrane protein, VF = virulence factor

**Table 3.** Top ten abundant proteins identified in the mono- and dual-species planktonic cultures of *F. nucleatum* and *P. gingivalis*.

| UniProt<br>Accession | Gene Name | Protein Description | Major Function | Log<sub>2</sub><br>LFQ |
| --- | --- | --- | --- | --- |
| Top 10 in *F. nucleatum* |  |  |  |  |
| Q8RGB0 | *FN0396* | Dipeptide-binding protein | peptide transporter | 32.37 |
| Q8REE2 | *FN1165* | D-galactose-binding protein | metal ion binding protein | 32.07 |
| Q8R643 | *FN1170* | Pyruvate-flavodoxin oxidoreductase | oxidoreductase | 31.77 |
| Q8RIP5 | *FN1526* | Fusobacterium outer membrane protein family | OMP/adhesin | 31.74 |
| Q8RHT4 | *FN1911* | Outer membrane protein | OMP | 31.44 |
| Q8REM0 | *FN1079* | Neutrophil-activating protein A | ferric iron binding protein | 31.42 |
| Q8RIN6 | *FN1535* | Acyl-CoA dehydrogenase, short-chain specific | oxidoreductase | 31.29 |
| Q8RG30 | *FN0488* | Glutamate dehydrogenase | oxidoreductase | 31.26 |
| Q8RGM9 | *FN0262* | Formate acetyltransferase | acetyltransferase | 31.25 |
| Q8RIM6 | *FN1549* | Stomatin like protein | --- | 31.13 |
| Top 10 in *P. gingivalis* |  |  |  |  |
| B2RHG7 | *ragA* | Receptor antigen A | protein with receptor activity | 33.52 |
| B2RHG8 | *ragB* | Receptor antigen B | protein with receptor activity | 32.97 |
| B2RM93 | *rgpA* | Arginine-specific cysteine proteinase RgpA | protease/VF | 32.93 |
| B2RLK7 | *hagA* | Hemagglutinin protein HagA | agglutinate erythrocytes/ VF | 32.66 |
| B2RLK2 | *kgp* | Lys-gingipain | protease/VF | 32.28 |
| B2RIQ1 | *PGN_0727* | 4-hydroxybutyryl-CoA dehydratase | oxidoreductase | 30.75 |
| B2RJ72 | *PGN_0898* | Probable peptidylarginine deiminase | deiminase/VF | 30.49 |
| B2RIQ3 | *PGN_0729* | Outer membrane protein 41 | OMP/ immunoreactive protein | 30.37 |
| B2RII3 | *PGN_0659* | 35 kDa hemin binding protein | hemin binding protein /VF | 30.02 |
| B2RKJ1 | *gdh* | NAD-specific glutamate dehydrogenase | oxidoreductase | 29.83 |
| Top 10 in dual-species planktonic culture |  |  |  |  |
| B2RHG7 | *ragA* | Receptor antigen A | protein with receptor activity | 33.14 |
| B2RHG8 | *ragB* | Receptor antigen B | protein with receptor activity | 32.92 |
| Q8RHY1 | *FN1859* | Major outer membrane protein FomA | OMP/VF | 32.39 |
| B2RM93 | *rgpA* | Arginine-specific cysteine proteinase RgpA | protease/VF | 31.91 |
| Q8RIP5 | *FN1526* | Fusobacterium outer membrane protein family | OMP/adhesin | 31.83 |
| B2RLK7 | *hagA* | Hemagglutinin protein HagA | agglutinate erythrocytes/ VF | 31.74 |
| Q8RG30 | *FN0488* | Glutamate dehydrogenase | oxidoreductase | 31.66 |
| Q8REE2 | *FN1165* | D-galactose-binding protein | metal ion binding protein | 31.61 |
| Q8R643 | *FN1170* | Pyruvate-flavodoxin oxidoreductase | oxidoreductase | 31.53 |
| Q8RGB0 | *FN0396* | Dipeptide-binding protein | peptide transporter | 31.50 |

Abbreviations: OMP = outer membrane protein, VF = virulence factor

Five of ten most abundant *F. nucleatum* and *P. gingivalis* proteins detected in the planktonic culture were also identified as the most abundant in the biofilm (**Table 2** and **3**). Yet, none of these proteins were among significantly changed proteins (see below) between the planktonic and biofilm conditions. Identification of the same proteins in both culturing conditions is likely a result of using the same medium and the same length of time for both planktonic culture and biofilm. *F. nucleatum* glutamate dehydrogenase (FN0488) is an example of such abundant protein; this protein is involved in oxidation-reduction process and can be used as a diagnostic marker for the genus *Fusobacterium* (34). Other *F. nucleatum* proteins detected at high levels were oxidoreductases, outer membrane proteins, and adhesins. Examples of these are FN1526 (known as RadD), which is an arginine-inhibitable adhesin required for inter-species adherence, and FN1859 (known as FomA), which functions as a non-specific porin and a virulence factor facilitating bacterial evasion from host immune surveillance (35). *P. gingivalis* proteins detected at high levels in planktonic culture and biofilm included virulence-related proteases cysteine proteinase RgpA and Lys-gingipain (Kgp). These proteins are involved in subversion of leukocytes and microbial dysbiosis, facilitating *P. gingivalis* colonization and the outgrowth of the surrounding microbial community (36).

To identify proteins produced in differential amounts by cells either in the biofilm or in planktonic culture, we statistically compared LFQ intensity readings by the student *t-test* (p≤0.05). Similarly, we compared the LFQ intensities of proteins produced by cells grown under either mono- or dual-species conditions (**Table 4**).

**Table 4.** Numbers of significantly changed proteins between different growth conditions.

| Culturing condition | Species | Proteins<sup>#</sup> | Significantly changed<sup>*</sup> |
| --- | --- | --- | --- |
| Bio vs Pla | Fn | 1070 | 5 |
| Bio vs Pla | Pg | 593 | 40 |
| Bio vs Pla | Fn/Pg | 797 | 112 |
| Bio | Fn vs Fn/Pg | 635 | 11 |
| Pla | Fn vs Fn/Pg | 998 | 10 |
| Bio | Pg vs Fn/Pg | 283 | 56 |
| Pla | Pg vs Fn/Pg | 242 | 63 |

<sup>#</sup> Number of quantified proteins used in two-sided t-test ( $p \leq 0.05$ ).
<sup>\*</sup> Proteins that passed the t-test and showed more than 2 $\log_2$ difference in the LFQ intensity.
Abbreviations: Fn - *F. nucleatum*, Pg - *P. gingivalis*, Fn/Pg – dual-species model, Bio - Biofilm, Pla = Planktonic culture

### F. nucleatum proteome is relatively similar under biofilm and planktonic conditions

Five out of 1,070 *F. nucleatum* proteins, which were quantified under both biofilm and planktonic culture, showed a significant change in their LFQ levels (**Fig 1A**). Proteins with significantly increased levels in the biofilm (**Table S4**) were Thiazole synthase (ThiC) and phosphomethylpyrimidine synthase (ThiG) involved in vitamin B1 (thiamine) and vitamin B2 (riboflavin) metabolic processes. It is currently unknown if vitamin B1 or B2 are aiding *F. nucleatum* biofilm formation. However, cells of the anaerobic gram-negative bacterium *Thermotoga maritima* that were grown as a biofilm exhibited increased transcription of genes involved in the biosynthesis of thiamine (37). *F. nucleatum* proteins with increased LFQ levels in the planktonic condition were a membrane transport protein (IIC component protein of the PTS system) and two transferases (Acetate CoA-transferase YdiF and Acetoacetate: butyrate/acetate coenzyme A transferase).

**Fig 1.**
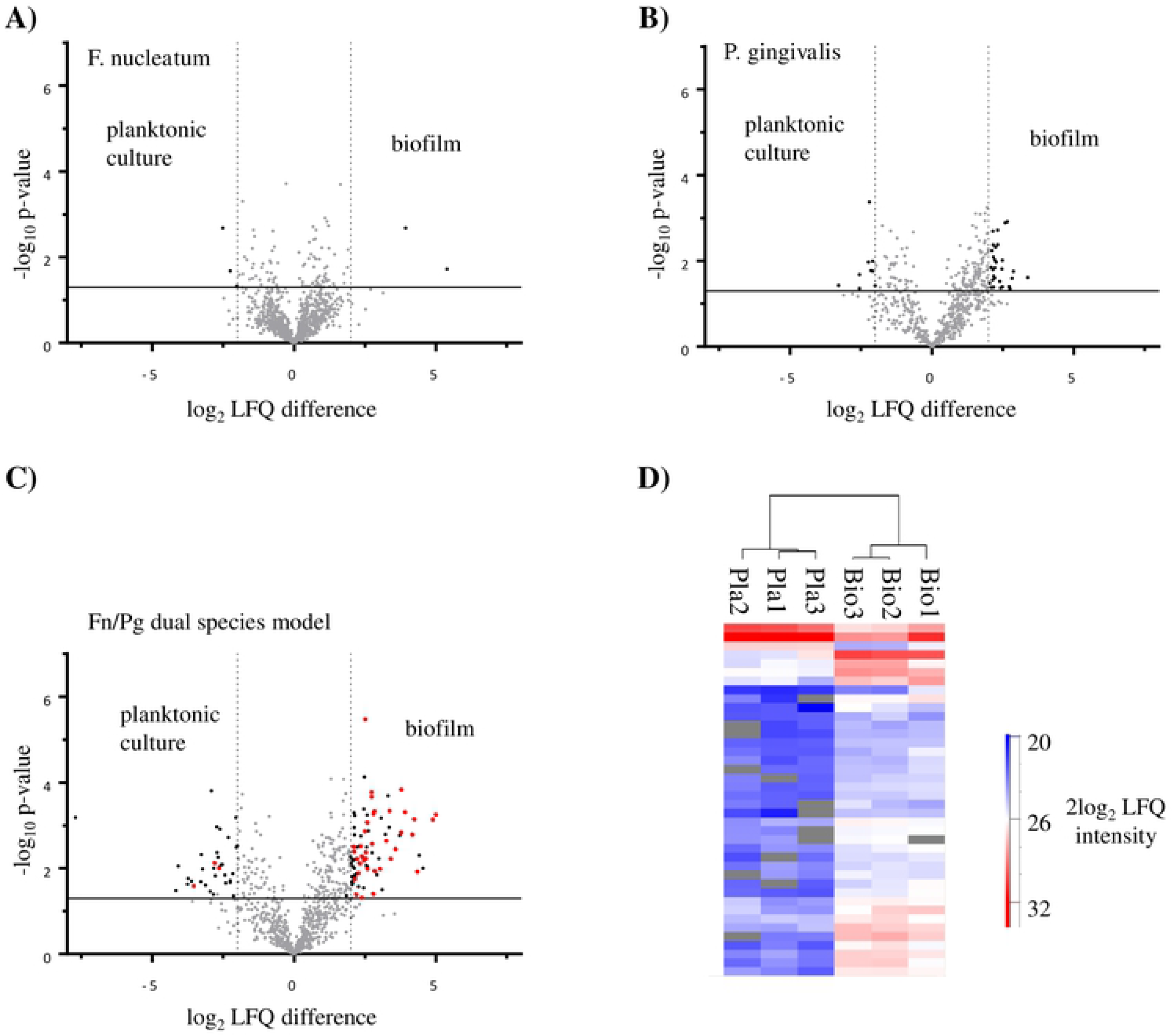
Differentially produced proteins in the biofilm and planktonic modes of growth. Volcano plots show results of t-test (p-value ≤0.05), which was performed on log2 LFQ intensities of proteins derived either from biofilm or planktonic cells of A) *F. nucleatum* (1069 proteins), B) *P. gingivalis* (593 proteins), and C) dual-species model (797 proteins). The t-test identified 5, 40, and 112 proteins as having significantly different levels (≥ 2log2 LFQ intensity) between the two conditions in A, B, and C, respectively (shown in black). The horizontal line shows the p-value cut-off of 0.05, and the vertical lines mark the log2 LFQ intensity = ±2. Red dots in C denote *P. gingivalis* proteins. D) Hierarchical clustering of 40 *P. gingivalis* proteins with significantly different log_2_ LFQ intensities in the dual-species model. Abbreviations: Pla – planktonic culture, Bio – biofilm. Biological replicates are shown, and grey fields indicate missing values, *i.e*., the protein was not detected, or its amounts were below quantifiable levels.

Some of the proteins associated with *F. nucleatum* pathogenicity were identified as significantly different (p ≤0.05) but showed less than 2log_2_ difference between the biofilm and planktonic cultures (**Table S4**). For example, metal-dependent hydrolase (FN1210), a resistance causing and drug efflux protein with beta-lactamase activity (38), had slightly increased LFQ levels in the biofilm mode of growth. Three other proteins (FN1613, FN0268, FN0235), which are virulence factors with peptidase activity and are involved in proteolysis, had also increased levels in the biofilm.

### P. gingivalis increases the production of some proteins when cultured in biofilm compared to the planktonic condition

Approximately 7% of all *P. gingivalis* proteins quantified under biofilm and planktonic culture (40 out of 593) showed significant changes in their LFQ levels (**Table S5**). We detected 30 proteins with more than 2log_2_ LFQ levels increase in the biofilm setting (**Fig 1B**). As in *F. nucleatum* biofilm, riboflavin biosynthesis protein (RibBA) had increased levels in the *P. gingivalis* biofilm. The latter coincides with a transcriptomic study, which showed that the protein is upregulated in a *P. gingivalis* biofilm (39). This protein is involved in biofilm-related functions, including quorum sensing signalling and extracellular electron transfer in different bacterial species (40, 41). Other proteins that increased in the biofilm were functional in translation (PG_RS01735, PG_RS01750, PG_RS08510, PG_RS05710) and amino acid biosynthesis (GpmA, PGN_0692).

We identified ten *P. gingivalis* proteins with increased amounts in the planktonic condition, including outer membrane efflux protein (PGN_1432) that has cellular transport activity, an integral component of membrane (PGN_0296), immunoreactive 23 kDa antigen protein (PGN_0482), and membrane-associated zinc metalloprotease (PGN_1582) that has peptidase activity and is involved in proteolysis. These results support findings from a gene expression analysis study, which showed that *P. gingivalis* genes are upregulated in the biofilm setting compared to planktonic culture (39).

### A majority of P. gingivalis proteins have increased levels in biofilm compared to planktonic culture when cultured as dual-species model with F. nucleatum

In the mixed-species cultures, 797 proteins were quantified under both biofilm and planktonic conditions, and LFQ intensities of 14% (112) proteins were significantly changed (**Fig 1C**). Among these proteins, we detected more proteins derived from *F. nucleatum* (72) than *P. gingivalis* (40). Of 78 proteins with increased amounts in the biofilm (**Table S6**), 41 derived from *F. nucleatum* and included the following functional clusters: lyase (9 proteins), metal binding (8 proteins), and energy production and conversion (4 proteins). The remaining 37 proteins were derived from *P. gingivalis* **(Fig 1D)** and included proteins with oxidoreductase (4 proteins) and translation (4 proteins) activity. These results are in accordance with findings from *P. gingivalis* mono-species culture, which also showed more proteins with increased levels in biofilm compared to the planktonic condition. In biofilm, *P. gingivalis* appeared to generally increase the production of proteins functional in translation, oxidation-reduction, and biosynthesis of riboflavin and amino acids. The identification and quantification of these proteins provide new insights into the physiology of this periodontal pathogen, adding additional information to previous gene expression studies (39, 42).

Most of the proteins with increased levels in the dual-species planktonic culture were derived from *F. nucleatum* (31 out of 34) and included 7 ribosomal proteins, 4 proteins involved in rRNA binding, and 6 translational proteins. Interestingly, the FadA adhesion proteins (FN0249 and FN0264) displayed almost 8-fold and 4-fold reduction, respectively, in the dual-species biofilm compared to the planktonic culture (**Table S6**), while showed no change in the mono-species cultures (**Table S4**). FadA helps *F. nucleatum* to adhere and invade host epithelial and endothelial cells (43). It also promotes colorectal carcinogenesis in humans by modulating signalling of E-cadherin/b-catenin, which was identified as the endothelial receptor for FadA (44). The reduction in FadA levels might be explained by a recent study, which showed that *P. gingivalis* could suppress an invasion of *F. nucleatum* into gingival epithelial cells (45). The authors attributed this to the degradation of E-cadherin by *P. gingivalis* gingipains (46). Our results confirm that the presence of *P. gingivalis* has a negative effect on *F. nucleatum* FadA proteins.

### F. nucleatum marginally changes its proteome in response to P. gingivalis presence

Eleven out of 635 *F. nucleatum* proteins that were quantified both in the mono- and dual-species biofilm showed significantly different levels (**Fig 2A** and **Table S7**). The three proteins with increased levels in the dual-species biofilm were an uncharacterized membrane protein (FN0514), transcriptional regulatory protein (FN0198), and multi-functional protein HppA that is involved in potassium ion transport. Proteins with increased levels in the mono-species biofilm included outer membrane proteins (FN0335, FN2103, and FN1449) and several uncharacterized proteins.

**Fig 2.**
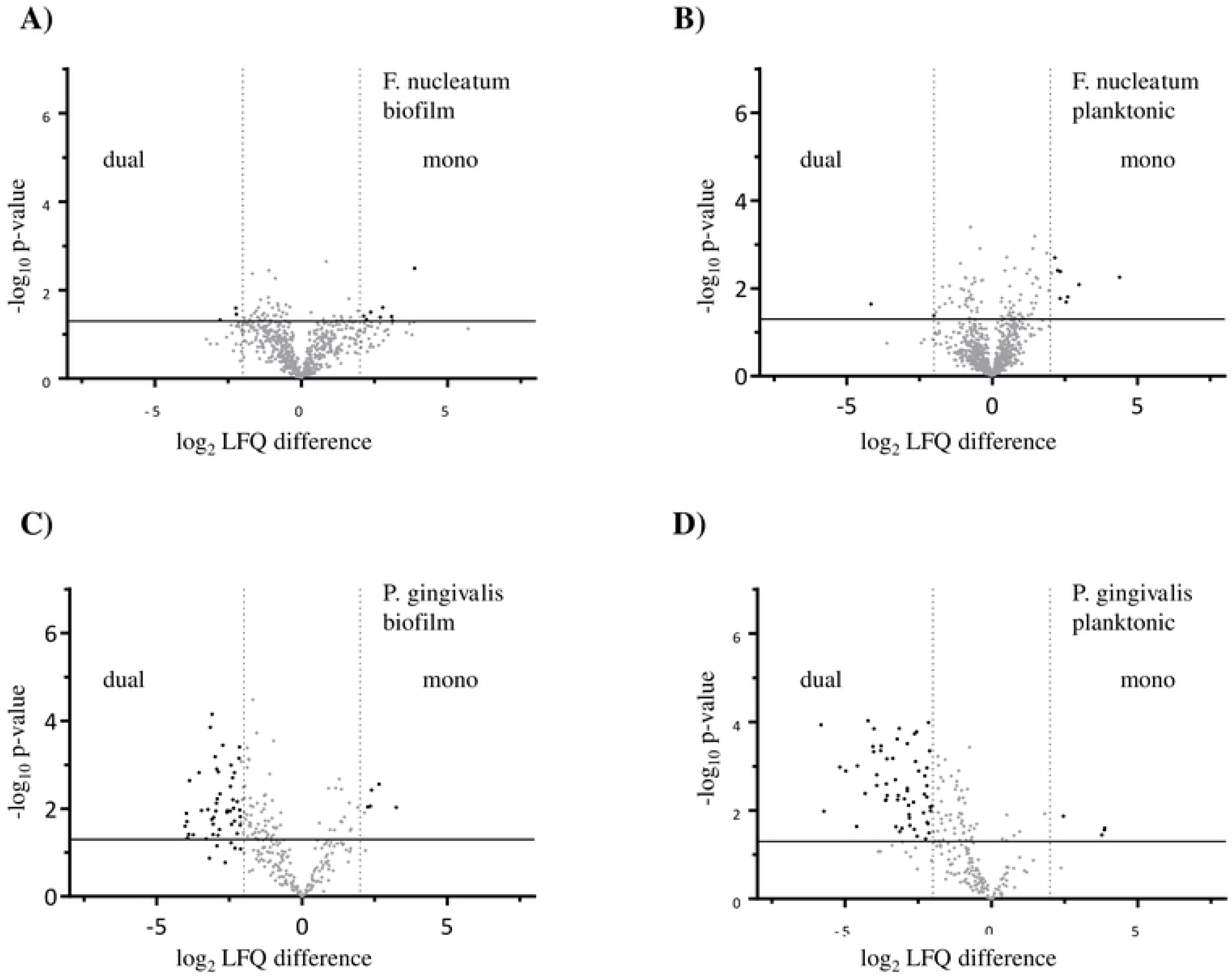
Differentially expressed proteins in mono- and dual-species biofilms and planktonic growth conditions. Volcano plot shows results of *t-test* with p-value ≤0.05, which was performed on log_2_ LFQ intensities for A) *F. nucleatum* biofilm vs dual-species biofilm (635 proteins), B) F. nucleatum planktonic vs dual-species planktonic (998 proteins), C) *P. gingivalis* biofilm vs dual-species biofilm (283 proteins), D) *P. gingivalis* planktonic vs dual-species planktonic (242 proteins). The t-test identified 11, 10, 56, and 63 proteins as having significantly different levels (≥ 2log_2_ LFQ intensity) between the two conditions in A, B, C, and D, respectively (shown in black). The horizontal line shows the p-value cut-off of 0.05, and the vertical lines mark the log_2_ LFQ intensity = ±2.

In the planktonic conditions, 10 out of 998 proteins showed significant changes. Two proteins had increased levels in the dual-species model, while 8 decreased in the mono-species culture (**Fig 2B** and **Table S8**). Most of these proteins were annotated as uncharacterized. Thus, the results suggest that *F. nucleatum* responds to the presence of *P. gingivalis* only mildly, mostly by decreasing production of specific proteins.

### P. gingivalis reduces its protein production in biofilm in response to F. nucleatum presence

We identified 283 *P. gingivalis* proteins that were quantified both in the mono- and dual-species biofilms, and most of the proteins with significantly different levels were decreased in the dual-species biofilm (51 out of 56) (**Fig 2C** and **Table S9**). Functional analysis of these proteins pointed out the following functional clusters: structural ribosomal activity (6 proteins), translation (6 proteins), oxidation-reduction (4 proteins), and RNA binding (4 proteins). The reduction in amounts of multiple proteins agreed with a proteomic study of *P. gingivalis*, where it was cultivated in three species community with *F. nucleatum* and *Streptococcus gordoni* (30). The study authors suggested that the microbial community provided physiologic support to *P. gingivalis* and, in this way, reduced its stress.

Statistical comparison of *P. gingivalis* proteins LFQ intensities between the mono-species and dual-species planktonic cultures identified 63 significantly different proteins, and 59 of these showed reduced levels in the dual-species culture (**Fig 2D** and **Table S10**). Twenty-six of the proteins with decreased levels in dual-species planktonic culture were also decreased in the dual-species biofilm condition (**Table S9**). In summary, when *P. gingivalis* cells were grown in the dual-species model, we detected a relatively high number of proteins that had reduced levels both in biofilm and planktonic culture. A plausible explanation for these findings, which are in line with previous studies (10, 11, 30), is that the presence of *F. nucleatum* proteins creates a supportive environment for *P. gingivalis* growth.

## Conclusion

This study has explored the expressed proteomes of two important oral pathogens and showed how different growth conditions affect their proteins’ qualitative and quantitative composition. While *F. nucleatum* proteome remained relatively similar under biofilm and planktonic conditions, *P. gingivalis* increased the production of a number of proteins when cultured in biofilm. The latter finding might reflect an adaptation of *P. gingivalis* to the biofilm condition. We identified the largest number of proteins (14%) with changed levels when the two species were grown together. These results show that the two bacteria actively interact with each other on the protein level and *F. nucleatum* presence influences the levels of multiple *P. gingivalis* proteins both in biofilm and planktonic culture. *P. gingivalis* showed greater protein changes compared to *F. nucleatum* in both settings (biofilm vs planktonic and mono-species vs dual-species model). The data support the notion that *P. gingivalis* adapts to the biofilm condition by increasing levels of certain proteins; however, the presence of *F. nucleatum* mitigates this need by providing more favourable growth conditions. Proteomic studies of multispecies biofilms are an important area for future investigations, and in combination with gingival cell culture models, such studies will provide important insights into the biofilm formation and, consequently, for the prevention of periodontal diseases.

## Author contributions

Study design (MMAM, VB, AHN, HGW), Funding (MMAM, VB), Experimental procedures (MMAM), Data analysis and interpretation (MMAM, VKP), Manuscript draft (MMAM, VKP), Text critical review (MMAM, VKP, AHN, HGW, VB).

## Acknowledgments

We acknowledge the Proteomic Unit at the University of Bergen, particularly Olav Mjaavatten, for the LC-MS/MS experiments support. This work was funded by grants from the Research Council of Norway and Gades Legat.

## References

1. Socransky SS, Haffajee AD, Cugini MA, Smith C, Kent RL, Jr. Microbial complexes in subgingival plaque. J Clin Periodontol. 1998;25(2):134–44.

2. Schaudinn C, Gorur A, Keller D, Sedghizadeh PP, Costerton JW. Periodontitis: an archetypical biofilm disease. J Am Dent Assoc. 2009;140(8):978–86.

3. Pihlstrom BL, Michalowicz BS, Johnson NW. Periodontal diseases. Lancet. 2005;366(9499):1809–20.

4. Kolenbrander PE. Oral microbial communities: biofilms, interactions, and genetic systems. Annu Rev Microbiol. 2000;54:413–37.

5. Holt SC, Ebersole JL. *Porphyromonas gingivalis, Treponema denticola,* and *Tannerella forsythia*: the “red complex”, a prototype polybacterial pathogenic consortium in periodontitis. Periodontol 2000. 2005;38:72–122.

6. Lamont RJ, Jenkinson HF. Life below the gum line: pathogenic mechanisms of Porphyromonas gingivalis. Microbiology and molecular biology reviews: MMBR. 1998;62(4):1244–63.

7. Naito M, Hirakawa H, Yamashita A, Ohara N, Shoji M, Yukitake H, et al. Determination of the genome sequence of Porphyromonas gingivalis strain ATCC 33277 and genomic comparison with strain W83 revealed extensive genome rearrangements in P. gingivalis. DNA Res. 2008;15(4):215–25.

8. Mysak J, Podzimek S, Sommerova P, Lyuya-Mi Y, Bartova J, Janatova T, et al. Porphyromonas gingivalis: major periodontopathic pathogen overview. Journal of immunology research. 2014;2014:476068.

9. Periasamy S, Kolenbrander PE. Mutualistic biofilm communities develop with Porphyromonas gingivalis and initial, early, and late colonizers of enamel. J Bacteriol. 2009;191(22):6804–11.

10. Metzger Z, Lin YY, Dimeo F, Ambrose WW, Trope M, Arnold RR. Synergistic pathogenicity of Porphyromonas gingivalis and Fusobacterium nucleatum in the mouse subcutaneous chamber model. J Endod. 2009;35(1):86–94.

11. Diaz PI, Zilm PS, Rogers AH. *Fusobacterium nucleatum* supports the growth of *Porphyromonas gingivalis* in oxygenated and carbon-dioxide-depleted environments. Microbiology. 2002;148(Pt 2):467–72.

12. Bradshaw DJ, Marsh PD, Watson GK, Allison C. Role of Fusobacterium nucleatum and coaggregation in anaerobe survival in planktonic and biofilm oral microbial communities during aeration. Infect Immun. 1998;66(10):4729–32.

13. Saito Y, Fujii R, Nakagawa KI, Kuramitsu HK, Okuda K, Ishihara K. Stimulation of Fusobacterium nucleatum biofilm formation by Porphyromonas gingivalis. Oral microbiology and immunology. 2008;23(1):1–6.

14. Polak D, Wilensky A, Shapira L, Halabi A, Goldstein D, Weiss EI, et al. Mouse model of experimental periodontitis induced by Porphyromonas gingivalis/Fusobacterium nucleatum infection: bone loss and host response. Journal of clinical periodontology. 2009;36(5):406–10.

15. Sauer K, Camper AK, Ehrlich GD, Costerton JW, Davies DG. Pseudomonas aeruginosa displays multiple phenotypes during development as a biofilm. J Bacteriol. 2002;184(4):1140–54.

16. Costerton JW. Introduction to biofilm. Int J Antimicrob Agents. 1999;11(3–4):217–21; discussion 37–9.

17. Llama-Palacios A, Potupa O, Sanchez MC, Figuero E, Herrera D, Sanz M. Proteomic analysis of Fusobacterium nucleatum growth in biofilm versus planktonic state. Mol Oral Microbiol. 2020;35(4):168–80.

18. Huang R, Li M, Gregory RL. Bacterial interactions in dental biofilm. Virulence. 2011;2(5):435–44.

19. Wisniewski JR, Zougman A, Nagaraj N, Mann M. Universal sample preparation method for proteome analysis. Nat Methods. 2009;6(5):359–62.

20. Rappsilber J, Mann M, Ishihama Y. Protocol for micro-purification, enrichment, pre-fractionation and storage of peptides for proteomics using StageTips. Nat Protoc. 2007;2(8):1896–906.

21. Cox J, Mann M. MaxQuant enables high peptide identification rates, individualized p.p.b.- range mass accuracies and proteome-wide protein quantification. Nat Biotechnol. 2008;26(12):1367–72.

22. Cox J, Hein MY, Luber CA, Paron I, Nagaraj N, Mann M. Accurate proteome-wide label-free quantification by delayed normalization and maximal peptide ratio extraction, termed MaxLFQ. Mol Cell Proteomics. 2014;13(9):2513–26.

23. Tyanova S, Temu T, Sinitcyn P, Carlson A, Hein MY, Geiger T, et al. The Perseus computational platform for comprehensive analysis of (prote)omics data. Nat Methods. 2016;13(9):731–40.

24. Huang da W, Sherman BT, Lempicki RA. Systematic and integrative analysis of large gene lists using DAVID bioinformatics resources. Nat Protoc. 2009;4(1):44–57.

25. Binns D, Dimmer E, Huntley R, Barrell D, O’Donovan C, Apweiler R. QuickGO: a web-based tool for Gene Ontology searching. Bioinformatics. 2009;25(22):3045–6.

26. Imai K, Asakawa N, Tsuji T, Akazawa F, Ino A, Sonoyama M, et al. SOSUI-GramN: high performance prediction for sub-cellular localization of proteins in gram-negative bacteria. Bioinformation. 2008;2(9):417–21.

27. Deutsch EW, Csordas A, Sun Z, Jarnuczak A, Perez-Riverol Y, Ternent T, et al. The ProteomeXchange consortium in 2017: supporting the cultural change in proteomics public data deposition. Nucleic Acids Res. 2017;45(D1):D1100–D6.

28. Jarnuczak AF, Vizcaino JA. Using the PRIDE Database and ProteomeXchange for Submitting and Accessing Public Proteomics Datasets. Curr Protoc Bioinformatics. 2017;59:13 31 1–13 31 12.

29. Xia Q, Wang T, Taub F, Park Y, Capestany CA, Lamont RJ, et al. Quantitative proteomics of intracellular Porphyromonas gingivalis. Proteomics. 2007;7(23):4323–37.

30. Kuboniwa M, Hendrickson EL, Xia Q, Wang T, Xie H, Hackett M, et al. Proteomics of Porphyromonas gingivalis within a model oral microbial community. BMC Microbiol. 2009;9:98.

31. Hendrickson EL, Wang T, Beck DA, Dickinson BC, Wright CJ, R JL, et al. Proteomics of Fusobacterium nucleatum within a model developing oral microbial community. Microbiologyopen. 2014;3(5):729–51.

32. Mohammed MM, Pettersen VK, Nerland AH, Wiker HG, Bakken V. Quantitative proteomic analysis of extracellular matrix extracted from mono- and dual-species biofilms of Fusobacterium nucleatum and Porphyromonas gingivalis. Anaerobe. 2017;44:133–42.

33. Bayles KW. The biological role of death and lysis in biofilm development. Nature Reviews Microbiology. 2007;5(9):721–6.

34. Gharbia SE, Shah HN. Characteristics of glutamate dehydrogenase, a new diagnostic marker for the genus Fusobacterium. J Gen Microbiol. 1988;134(2):327–32.

35. Toussi DN, Liu X, Massari P. The FomA porin from Fusobacterium nucleatum is a Toll-like receptor 2 agonist with immune adjuvant activity. Clin Vaccine Immunol. 2012;19(7):1093–101.

36. Zenobia C, Hajishengallis G. Porphyromonas gingivalis virulence factors involved in subversion of leukocytes and microbial dysbiosis. Virulence. 2015;6(3):236–43.

37. Pysz MA, Conners SB, Montero CI, Shockley KR, Johnson MR, Ward DE, et al. Transcriptional analysis of biofilm formation processes in the anaerobic, hyperthermophilic bacterium Thermotoga maritima. Appl Environ Microbiol. 2004;70(10):6098–112.

38. Kumar A, Thotakura PL, Tiwary BK, Krishna R. Target identification in Fusobacterium nucleatum by subtractive genomics approach and enrichment analysis of host-pathogen protein-protein interactions. BMC Microbiol. 2016;16:84.

39. Romero-Lastra P, Sanchez MC, Ribeiro-Vidal H, Llama-Palacios A, Figuero E, Herrera D, et al. Comparative gene expression analysis of Porphyromonas gingivalis ATCC 33277 in planktonic and biofilms states. PLoS One. 2017;12(4):e0174669.

40. Rajamani S, Bauer WD, Robinson JB, Farrow JM, 3rd, Pesci EC, Teplitski M, et al. The vitamin riboflavin and its derivative lumichrome activate the LasR bacterial quorum-sensing receptor. Mol Plant Microbe Interact. 2008;21(9):1184–92.

41. Marsili E, Baron DB, Shikhare ID, Coursolle D, Gralnick JA, Bond DR. Shewanella secretes flavins that mediate extracellular electron transfer. Proc Natl Acad Sci U S A. 2008;105(10):3968–73.

42. Romero-Lastra P, Sanchez MC, Llama-Palacios A, Figuero E, Herrera D, Sanz M. Gene expression of Porphyromonas gingivalis ATCC 33277 when growing in an in vitro multispecies biofilm. PLoS One. 2019;14(8):e0221234.

43. Fardini Y, Wang X, Temoin S, Nithianantham S, Lee D, Shoham M, et al. Fusobacterium nucleatum adhesin FadA binds vascular endothelial cadherin and alters endothelial integrity. Mol Microbiol. 2011;82(6):1468–80.

44. Rubinstein MR, Wang X, Liu W, Hao Y, Cai G, Han YW. Fusobacterium nucleatum promotes colorectal carcinogenesis by modulating E-cadherin/beta-catenin signaling via its FadA adhesin. Cell Host Microbe. 2013;14(2):195–206.

45. Jung Y-J, Jun H-K, Choi B-K. Porphyromonas gingivalis suppresses invasion of Fusobacterium nucleatum into gingival epithelial cells. Journal of oral microbiology. 2017;9(1):1320193.

46. Katz J, Yang QB, Zhang P, Potempa J, Travis J, Michalek SM, et al. Hydrolysis of epithelial junctional proteins by Porphyromonas gingivalis gingipains. Infection and immunity. 2002;70(5):2512–8.

